# De Novo Design and Computational Validation of a High-Affinity Peptide Inhibitor Targeting the HPV E1-E2 Interface

**DOI:** 10.64898/2026.06.01.729313

**Authors:** Sean Fletcher, Esther E. Biswas-Fiss, Subhasis Biswas

## Abstract

The oncogenic progression of high-risk Human Papillomavirus (HPV) strains relies on the cooperative interaction between the E1 replicative helicase and the E2 origin-binding protein to initiate viral DNA amplification. Disrupting this protein-protein interaction represents a promising, yet clinically unrealized, therapeutic paradigm for treating established HPV infections prior to malignant transformation. This study presents a comprehensive computational pipeline for the de novo design and evaluation of peptide inhibitors targeting the HPV E1-E2 interface, specifically a conserved arginine triad on the solvent-exposed surface of the E1 helicase. AlphaProteo was used for sequence discovery, and AlphaFold 3 for complex structural prediction, generating a candidate library that was subsequently subjected to dual-scale Molecular Dynamics (MD) simulations and MM/GBSA thermodynamic validation using GROMACS. Binder 8 emerged as the lead candidate, yielding a predicted binding free energy of -59.1 ± 0.7 kcal/mol — a statistically significant improvement over the native E1-E2 baseline (Welch’s t-test, p = 8.14e-19; Cohen’s d = 2.21). As an implicit solvent method, MM/GBSA overestimates absolute affinities; reported values reflect effective binding enthalpy and should be interpreted as relative rankings. Per-residue energy decomposition confirms binding is anchored through multi-point interactions with the arginine triad. Physicochemical profiling via CSM-Toxin and AlgPred 2.0 confirms zero predicted toxicity and non-allergenic properties for Binder 8. Sequence alignment across 183 oncogenic Alpha-papillomavirus genotypes demonstrates near-universal conservation of the targeted triad, supporting Binder 8 as a candidate scaffold for broad-spectrum antiviral development. These findings provide a computationally validated blueprint for future in vitro validation via Bio-layer interferometry.

## 1. Introduction

### 1.1 The Epidemiological Burden of Human Papillomavirus

The clinical and epidemiological burden imposed by Human Papillomavirus (HPV) represents a major global public health challenge defined by significant geographic disparities. High-risk HPV (HR-HPV) genotypes, most prominently HPV-16 and HPV-18 are the definitive etiological agents for a broad and devastating spectrum of malignancies, including cervical, anogenital, and oropharyngeal squamous cell carcinomas. [1] Epidemiological data from recent global oncology assessments emphasizes the severity of this ubiquitous oncogenic pathogen. In 2022, cervical cancer alone accounted for 44.0% of all HPV-associated cancers globally, translating to approximately 662,044 newly diagnosed cases with an age-standardized incidence rate (ASIR) of 14.1 per 100,000 individuals. [1] Furthermore, HPV-driven head and neck cancers, specifically targeting the oropharynx, contributed to 45.5% of cases (685,204 incidents), exhibiting a demographic disparity where the ASIR in males (11.2 per 100,000) was nearly quadruple that observed in females (3.1 per 100,000). [1]

Additionally, the socioeconomic distribution of this oncogenic burden remains highly disproportionate. The highest cervical cancer incidence and mortality rates are concentrated in developing nations and low-resource settings, such as Eswatini (ASIR 77.4) and Zimbabwe (ASIR 60.4), contrasting with significantly lower rates in high-resource regions like Saudi Arabia and Kuwait. [1] While widespread vaccination campaigns utilizing bivalent, quadrivalent, and nonavalent vaccines have demonstrated remarkable efficacy in preventing primary infections, they are inherently restricted to a purely prophylactic role. [2] These vaccines function by eliciting neutralizing immunoglobulins against the L1 major capsid protein but offer no therapeutic benefit or viral clearance for individuals with established, persistent cellular infections. [2] In patient populations with compromised immune surveillance, such as individuals living with HIV, the incidence, persistence, and rapid progression of HR-HPV infections to cervical intraepithelial neoplasia (CIN) and invasive carcinomas are significantly amplified. Globally, the most aggressive of these high-risk strains, HPV-16 and HPV-18, account for roughly 76% of all cervical cancer diagnoses. [2] The complete absence of targeted antiviral therapeutics capable of clearing persistent episomal or integrated HR-HPV infections necessitates the discovery of novel pharmacological interventions that directly inhibit the viral replication machinery before oncogenic integration occurs.

### 1.2 Structural Biology of the E1 Helicase and the E1-E2 Protein Interaction

The replication of the compact, double-stranded HPV genome is entirely dependent on the cooperative, highly regulated function of two virus-encoded early proteins: the E1 replicative DNA helicase and the E2 regulatory and origin-binding protein. Because the papillomavirus genome is small and does not encode its own DNA polymerase, primase, or accessory replication factors, it must hijack the host cell’s DNA replication machinery. [3] The initiation of this episomal amplification process requires the precise assembly of the E1-E2 complex directly at the viral origin of replication. [4]

The structural biology underlying the E1-E2 interaction is characterized by a dynamically regulated protein-protein interface. The E1 protein is a highly conserved member of the Superfamily 3 AAA+ (ATPases Associated with various cellular Activities) helicase family. [5] Its structure comprises an N-terminal regulatory domain containing a nuclear localization signal, a central DNA-binding domain responsible for recognizing inverted-repeat sequences at the viral origin, and a large C-terminal helicase core. This helicase core contains classical Walker A (P-loop) and Walker B motifs for nucleotide binding and ATP hydrolysis, alongside a highly conserved arginine finger that extends into the catalytic pocket of adjacent subunits during hexamerization to stimulate ATP hydrolysis. [6]

The E2 protein functions as a molecular matchmaker during the initiation phase. [7] It consists of a largely alpha-helical N-terminal transactivation domain, a flexible hinge region, and a C-terminal beta-barrel DNA-binding domain that acts to tether the complex to specific target sequences. [3] Crystallographic studies of the HPV-18 E1-E2 complex (specifically structural models such as PDB: 1TUE) reveal that the E2 transactivation domain binds to the C-terminal helicase domain of E1, forming a distinctive, rigid “C”-shaped heterodimeric assembly. [7] The primary regulatory function of E2 is driven by direct steric occlusion. By binding to the E1 monomer, the C-terminal beta-strand domain of E2 physically shields an important oligomerization interface on E1, directly blocking approximately 10% of E1’s functional volumetric space. [7] This steric activity prevents E1 from prematurely assembling into the double-hexameric ring required for active DNA melting and unwinding. E2 essentially acts as a highly specific chaperone, guiding the E1 protein to the viral origin. Once successfully localized to the origin DNA, the binding of ATP acts as an allosteric effector, inducing a conformational shift that dramatically lowers the affinity between the two proteins, triggering the necessary dissociation of E2. Only upon E2 dissociation can E1 recruit additional identical subunits to form the active, functional hexameric helicase. [7] Therefore, the E1-E2 physical interaction is an absolute, non-negotiable prerequisite for viral replication, making it a highly attractive, yet historically challenging, target for pharmacological intervention.

### 1.3 The Rationale for Targeting the E1 Arginine Triad

Protein-protein interactions (PPIs) are notoriously difficult to target with traditional small-molecule therapeutics due to their expansive, topologically flat, and predominantly featureless hydrophobic interfaces. [8] However, structural and electrostatic analysis of the E1-E2 binding interface has revealed specific residues that mediate the high-affinity binding of the two viral proteins. A dominant feature of this exact interface on the solvent-exposed surface of the E1 helicase is a highly basic surface patch composed of three critical arginine residues within close proximity: R447, R454, and R622. [7]

This arginine triad serves as the primary electrostatic binding site for the complementary acidic residues embedded within the E2 transactivation domain. Previous mutagenesis studies have shown that disrupting this triad, notably through the R454A mutation, selectively eliminates the E1-E2 interaction. This cripples viral replication by preventing E1 recruitment to the origin, proving that the interaction interface can be targeted without destabilizing the intrinsic folding or isolated enzymatic activity of the E1 helicase. [7] The specific geometry and high positive charge density of these three arginine side chains make them an ideal structural target for a complementary binding strategy analogous to an arginine trap. Arginine traps are highly specialized, high-affinity binding mechanisms, frequently observed in the natural maturation of germline-encoded autoantibodies. [9] In an arginine trap, a highly complementary arrangement of acidic or sterically constrained amino acids forms a localized electronegative pocket that neutralizes the delocalized positive charge of the target guanidinium group. [9] By designing a de novo peptide inhibitor that precisely mimics the acidic engagement of the native E2 protein and explicitly traps the E1 arginine triad, it is theoretically possible to competitively and irreversibly occlude the E2 binding site. A biologic peptide rationally designed to exploit this specific structural vulnerability would actively prevent the formation of the E1-E2 pre-initiation complex, thereby halting viral DNA replication at its earliest possible stage.

### 1.4 Generative AI in Drug Discovery

The conventional pharmaceutical pipeline for discovering novel peptide inhibitors and biologic therapeutics relies heavily on strenuous, resource-intensive, and time-consuming empirical methodologies such as phage display, high-throughput biochemical screening, and iterative rounds of slow crystallographic optimization. [10–12] These historical processes suffer from high developmental attrition rates and immense financial costs.

However, the recent advent of generative artificial intelligence and deep-learning language models has fundamentally shifted this biological paradigm, enabling the precise, on-demand de novo design of high-affinity, target-specific protein binders. [13]

At the forefront of this computational advancement is AlphaProteo, a novel generative AI system developed by Google DeepMind specifically engineered to design novel, highly stable protein binders for structurally diverse target molecules. [14] Unlike purely predictive or analytical models, AlphaProteo is a generative framework capable of understanding the highly complex stereochemical, entropic, and electrostatic requirements of a designated target epitope, then generating entirely novel amino acid sequences that exhibit high geometric and energetic complementarity. Empirical wet-lab validations of the AlphaProteo architecture have demonstrated that it consistently achieves experimental hit rates and binding affinities that are 3 to 300 times superior to previous state-of-the-art computational methods, frequently generating de novo binders with sub-nanomolar to picomolar affinities in a single round of medium-throughput screening. [14] The system has already successfully generated neutralizing binders for highly complex targets, including VEGF-A, IL-7Rα, and the SARS-CoV-2 spike protein receptor-binding domain. [14]

Complementing this generative function is AlphaFold 3, which is capable of modeling structural interactions between proteins, DNA, RNA, simple ions, and small-molecule ligands with atomic-level precision, achieving a documented 50% improvement in structural accuracy over traditional physics-based docking methodologies on standard benchmarks like PoseBusters. [15,16] By integrating AlphaProteo for primary sequence generation and AlphaFold 3 for structural prediction of the complex, researchers can effectively bypass months or even years of costly empirical screening to immediately identify plausible therapeutic candidates ready for thermodynamic and cellular evaluation. **(Fig. 1)**

**Fig. 1.**
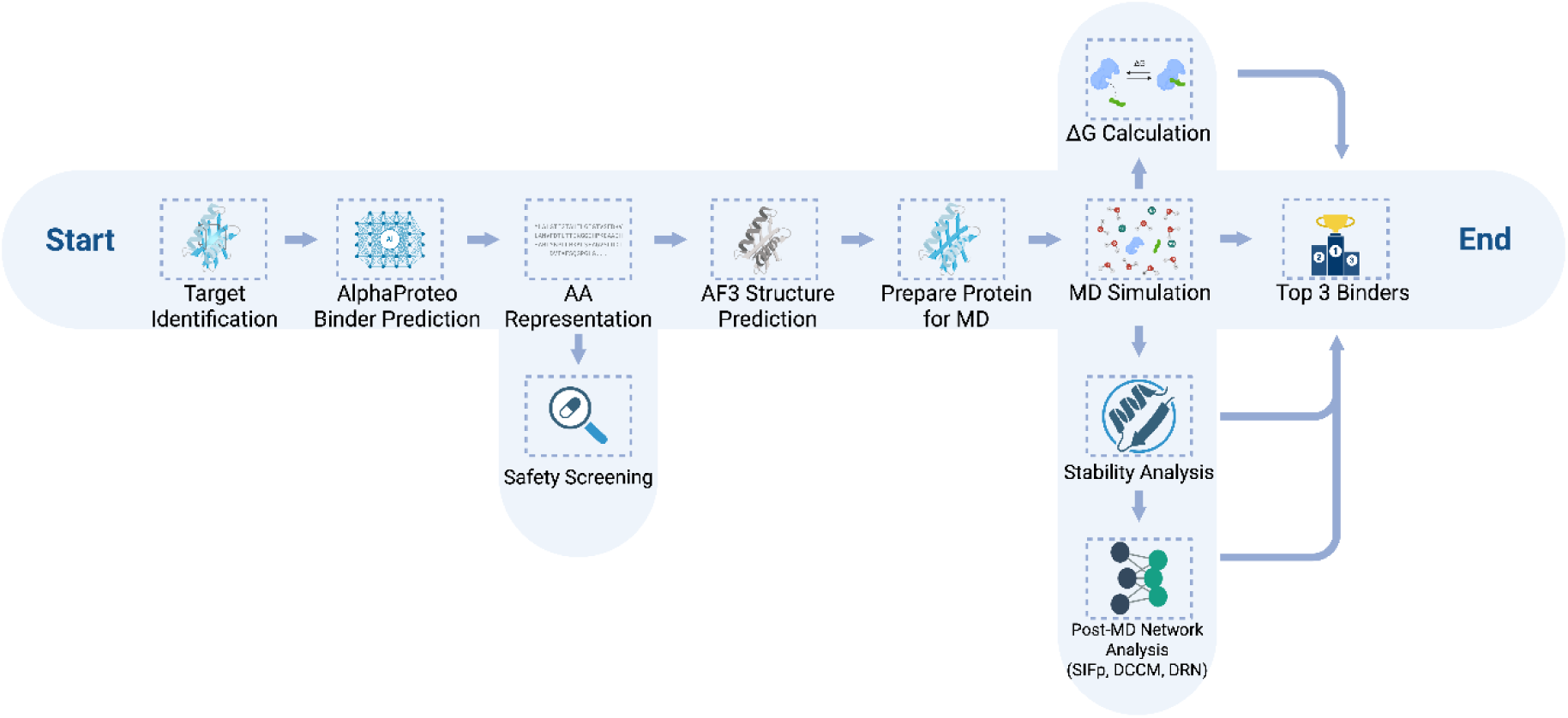
De novo generative design and validation framework targeting the HPV-18 E1 helicase. The computational pipeline initiates with target identification, focusing specifically on the conserved arginine triad of the E1 protein. The AlphaProteo generative AI model is employed to predict novel amino acid binder sequences. These sequences undergo preliminary safety screening to evaluate pharmacological properties like toxicity and allergenicity. AlphaFold 3 is then used to predict the three-dimensional structures of the resulting peptide-receptor complexes. Following preparation, the complexes are subjected to molecular dynamics (MD) simulation to capture biophysical behavior and structural relaxation. The simulation trajectories inform stability analysis, post-MD network analysis, and binding free energy calculations (ΔG). This comprehensive filtering process ultimately identifies the top three optimized, high-affinity binders.

## 2. Methods

### 2.1 Generative Design and Complex Prediction

The initial phase of the in silico biologic drug discovery pipeline utilized the AlphaProteo generative model to design a library of peptide binder sequences specifically targeting the solvent-exposed E1 helicase interface of the HPV-18 genotype. The target interface was defined around the geometric coordinates of the arginine triad (R447, R454, R622). An initial library of 14 candidate sequences was generated. **(Table S1)** AlphaFold 3 was subsequently utilized to generate three-dimensional models of the predicted Receptor-Peptide interactions for the initial candidate pool. To serve as an accurate and comparable baseline, the native HPV-18 E1-E2 complex (derived directly from the PDB sequence 1TUE) was also predicted using AlphaFold 3. [7] To systematically narrow this pool, a multi-tiered filtering protocol was employed. First-line structural filtering utilized an AlphaFold 3 interface predicted template metric (ipTM) threshold of ≥ 0.80 to eliminate low-confidence complexes. The surviving candidates were then visualized via PyMOL to verify direct interfacial contact with the target arginine triad residues across both HPV-18 and HPV-16 predicted models. This pre-screening isolated the top-ranking candidates for further evaluation.

### 2.2 In Silico Safety, Toxicity, and Physicochemical Profiling

Translating a computationally generated binder into a viable clinical biologic requires strict safety, immunogenicity, and stability profiling. To pre-emptively filter candidate sequences harboring toxic and/or immunogenic properties, a multi-tiered in silico screening protocol was implemented. Cellular toxicity was evaluated using the CSM-Toxin deep-learning server, utilizing a binary classification threshold of 0.968 to determine toxicity. CSM-Toxin is built upon the robust ProteinBERT architecture, a natural language processing-inspired model pre-trained on an exhaustive database of over 100 million sequences using Masked Language Modeling. [17] It achieves strong predictive accuracy by reading primary amino acid sequences as contextual features without requiring external, three-dimensional structural features. [18] Immunogenic and allergenic risks were quantified using the AlgPred 2.0 computational server, a predictive algorithm utilizing autocross covariance, sequence mapping, and machine learning to map known IgE epitopes and accurately predict whole-sequence allergenicity. [19]

Finally, the physicochemical stability, aqueous solubility, and estimated pharmacokinetic parameters (including the in vivo half-life and the instability index) were determined utilizing the ProtParam tool hosted on the ExPASy bioinformatics portal. [20] The Grand Average of Hydropathy (GRAVY) score was calculated to ensure the designs maintain high hydrophilicity, preventing undesirable aggregation in physiological human serum.

### 2.3 System Preparation and Molecular Dynamics Protocol

To accurately capture the biophysical behavior, structural relaxation, and interaction persistence of the predicted complexes, Molecular Dynamics (MD) simulations were executed utilizing the GROMACS v2025.4 simulation engine. [21,22] The predicted Receptor-Peptide complexes were parameterized utilizing the CHARMM36m all-atom force field, which is specifically optimized for protein folding dynamics and complex PPIs. [23] Given the highly basic nature of the targeted arginine triad, accurately capturing the local electrostatic environment was a critical consideration. Standard uncharged or capped peptide termini often lead to artificial fraying or unnatural repulsion during MD simulations. [24] Therefore, the designed peptides were explicitly modeled with highly charged zwitterionic termini, an NH3^+^ group at the N-terminus and a COO^-^ group at the C-terminus. This strict parameterization protocol provided the necessary electrostatic anchors to stabilize the peptide terminals against the substantial positive charge exerted by the surface arginines.

The protein systems were solvated within a cubic simulation box, maintaining a 1.0 nm minimum distance between the outermost solute atoms and the periodic box edges to prevent self-interaction artifacts across periodic boundaries. Solvation was achieved using the CHARMM-modified TIP3P explicit water model, replicating the dielectric constant, heat capacity, and kinetic diffusion properties of bulk physiological water. [25,26] The overall charge of each system was mathematically neutralized, and a physiological saline environment of 0.15 M NaCl was established through the targeted addition of sodium and chloride ions, mimicking the human intracellular environment.

Prior to integrating the equations of motion, structural relaxation was performed using a steepest descent energy minimization algorithm to resolve localized steric clashes generated during the AlphaFold 3 pose prediction. Energy minimization was performed until the maximum threshold force (F_max_) converged below 1000.0 kJ/mol/nm. The systems were then subjected to a two-stage equilibration process to establish correct thermodynamic ensembles without denaturing the proteins. Both equilibration phases were integrated using a continuous 1 fs time step. First, a 50 ps NVT (constant Number of particles, Volume, and Temperature) equilibration was conducted at 300 K utilizing the V-rescale thermostat (coupling time constant (τ_t_ = 0.1 ps) to stabilize the kinetic energy and uniformly distribute heat. [27] This was immediately followed by a 50 ps NPT (constant Number of particles, Pressure, and Temperature) equilibration at a standard atmospheric pressure of 1.0 bar using the C-rescale barostat (τ_p_ = 2.0 ps, isothermal compressibility = 4.5 × 10^-5^ bar^-1^) to ensure proper solvent density and box volume stabilization. [28]

The production MD simulations utilized a standard integration time step of 0.002 ps (2 fs). All covalent bonds involving fast-vibrating hydrogen atoms were rigidly constrained using the Linear Constraint Solver (LINCS) algorithm, allowing for the 2-fs step. [29] A non-bonded cutoff radius of 1.2 nm was enforced for both van der Waals and short-range electrostatic interactions, while long-range electrostatic forces were computed using the Particle Mesh Ewald mathematical method. [30] To ensure both long-term structural stability and high statistical confidence in the final thermodynamic scoring, a dual-scale simulation scheme was deployed. First, a high-throughput statistical ensemble phase was performed to determine the predicted binding affinities of each candidate and maximize the statistical power of the calculations. [31–34] This involved launching 50 independent, short-trajectory replicates per system across 6 systems (Binders 6, 8, 10, 11, 13, and 1TUE), yielding 300 total independent runs. [32] Each independent replicate was seeded with randomized initial Maxwell-Boltzmann velocities (gen_seed = -1) to explore diverse localized energetic microstates, followed by 100 ps of re-equilibration and 100 ps of production sampling. [35] This short-trajectory ensemble strategy directly adheres to the established framework of Genheden and Ryde [32], who demonstrated that running multiple independent replicas with brief production windows (20 to 200 ps) provides superior phase-space sampling and thermodynamic convergence compared to single long simulations. Based on the statistical ensemble data, the top 3 binders and the native E1-E2 complex (PDB: 1TUE) were isolated and advanced to the stability analysis phase. This phase involved subjecting the complexes to triplicate 50 ns continuous production runs, yielding 150 ns total sampling per system to confirm the persistence of the interfacial contacts and the avoidance of complex dissociation. [35] Structural metrics, including the Radius of Gyration (Rg) and Root Mean Square Deviation (RMSD), were smoothed utilizing a 150-frame rolling mean filter (representing a 1.5 ns window) to emphasize long-term stability trends. Per-residue Root Mean Square Fluctuations (RMSF) were similarly smoothed using a 5-frame rolling mean filter to maintain local resolution while reducing high-frequency vibrational noise. [35] Due to the N-terminal truncation of the E1 helicase domain used in the simulation models, the highly conserved biological Arginine Triad (R447, R454, and R622) corresponds to topological indices R20, R27, and R195, respectively. To align with the extracted thermodynamic and structural data, these targeted residues will hereafter be referred to as R20, R27, and R195.

### 2.4 Thermodynamic Scoring

The predicted binding free energies (ΔG_MM/GBSA_) for all 50 independent replicates were calculated using the Molecular Mechanics/Generalized Born Surface Area (MM/GBSA) method as implemented in the gmx_MMPBSA (v1.6.4) analytical Python suite. [34] MM/GBSA provides a highly effective, widely validated balance between computational efficiency and thermodynamic accuracy for evaluating relative binding affinities in large protein-protein complexes. [36] The thermodynamic calculations used the Onufriev-Bashford-Case (OBC2) Generalized Born implicit solvent model (igb = 5). [37] The implicit solvent environment was parameterized with a physiological salt concentration of 0.15 M, an internal solute dielectric constant of 1.0 (representing the hydrophobic protein core), and an external solvent dielectric constant of 80.0 to accurately represent the high polarizability of surrounding water. The non-polar solvation free energy, representing the hydrophobic effect, was calculated using a standard surface tension parameter of 0.0072 kcal/mol/Å^2^.

Furthermore, to achieve a better understanding of the binding architecture, mechanistic insights were extracted utilizing the per-residue energy decomposition module within gmx_MMPBSA (idecomp = 2), allowing for the analysis of individual side-chain and backbone enthalpic contributions to the global binding free energy. [36] All statistical margins of error reported in the results reflect the standard error of the mean (SEM) derived across the 50 independent replicates, ensuring a statistically rigorous estimation of the binding affinity devoid of single-run long-run anomalies. [31,32,38] To formally evaluate the significance of the thermodynamic differences between candidate binders and the native reference, standard deviations (SD) were calculated. Welch’s t-tests were performed on the replicate distributions with significance thresholds adjusted via Bonferroni correction for multiple comparisons (p < 0.0167). Additionally, Cohen’s d was calculated to quantify the biological effect size of the binding improvements.

### 2.5 Conformational Dynamics and Allosteric Network Analysis

To evaluate the large-scale allosteric effects of the de novo peptide inhibitors on the E1 protein, Dynamic Cross-Correlation Matrix (DCCM) and Dynamic Residue Network (DRN) analyses were executed across the triplicate production trajectories. These topological assessments provide insight into how the peptides alter the internal communication and conformational flexibility of the viral protein. DCCM was employed to quantify the correlated and anti-correlated motions between all alpha-carbon pairs during the trajectory. [39,40] To standardize cross-system comparison, given the differing system sizes (1TUE: ∼419 residues; binder complexes: ∼253 residues), DCCM pair counts at the |r| > 0.6 threshold were normalized as fractions of the total number of possible unique residue pairs (*N* (*N* - 1) / 2). Hydrogen bond analysis was restricted to interfacial contacts between the receptor and ligand chains using MDAnalysis, HydrogenBondAnalysis, excluding intra-chain H-bonds that scale proportionally with system size. The Radius of Gyration was calculated over the shared E1 receptor domain only (204 residues) to eliminate baseline size differences. Finally, to map the internal allosteric communication pathways within the inhibited complex, Betweenness Centrality (BC) scores were calculated on the shared 204-residue E1 receptor subgraph to ensure identical normalization across all systems. [41] The underlying graphs were constructed as a consensus residue network averaged over time and across replicates from every 10th trajectory frame. Following the MD-TASK framework, network edges were defined by a 6.7 Å non-hydrogen minimum distance cutoff and filtered using a strict 75% contact persistence threshold. The pre-established correlation threshold of |r| > 0.6 served to capture significant allosteric communication while filtering out background noise.

### 2.6 Protein Sequence Alignment

To evaluate the conservation of the targeted protein interface, full E1 protein sequences were acquired from the Papillomavirus Episteme (PaVE) database. [42,43] Two distinct datasets were compiled: a comprehensive library of the oncogenic Alpha-papillomavirus genus including reference sequences and clinical subtypes (n=183), and a broader evolutionary baseline encompassing reference types from the Alpha, Beta, and Gamma (ABG) genera (n=194). **(Table S2)** Multiple sequence alignments were performed utilizing the MAFFT algorithm configured with the E-INS-i strategy to accurately align sequences containing multiple conserved domains and extensive unalignable regions. [44] Following alignment, the target arginine triad (R20, R27, R195) was analyzed to quantify both strict sequence identity and physicochemical conservation, defined as the maintenance of a basic Arginine or Lysine residue at each respective position.

## 3. Results

### 3.1 Generative Design and Structural Prediction

Following the AI-driven generation of candidate sequences, the full three-dimensional conformations of the resulting Receptor-Peptide complexes were predicted using the AlphaFold 3 architecture. To establish an accurate baseline for comparison, the native HPV-18 E1-E2 complex (PDB: 1TUE) was also predicted. **(Fig. 2).** Structural alignment of this predicted baseline against the native crystal structure yielded an all-atom RMSD of 0.674 Å over 2523 aligned atoms. **(Fig. S1)** Structural filtering of the initial 14 candidates, using a strict ipTM threshold and analysis for electrostatic interactions between the arginine triad successfully eliminated low-confidence or non-interacting models (Binders 3, 7, 9, 12, and 14). PyMOL visualization for arginine triad binding of the remaining candidates isolated Binders 6, 8, 10, 11, and 13 as the primary high confidence leads for downstream MD analysis.

**Fig. 2.**
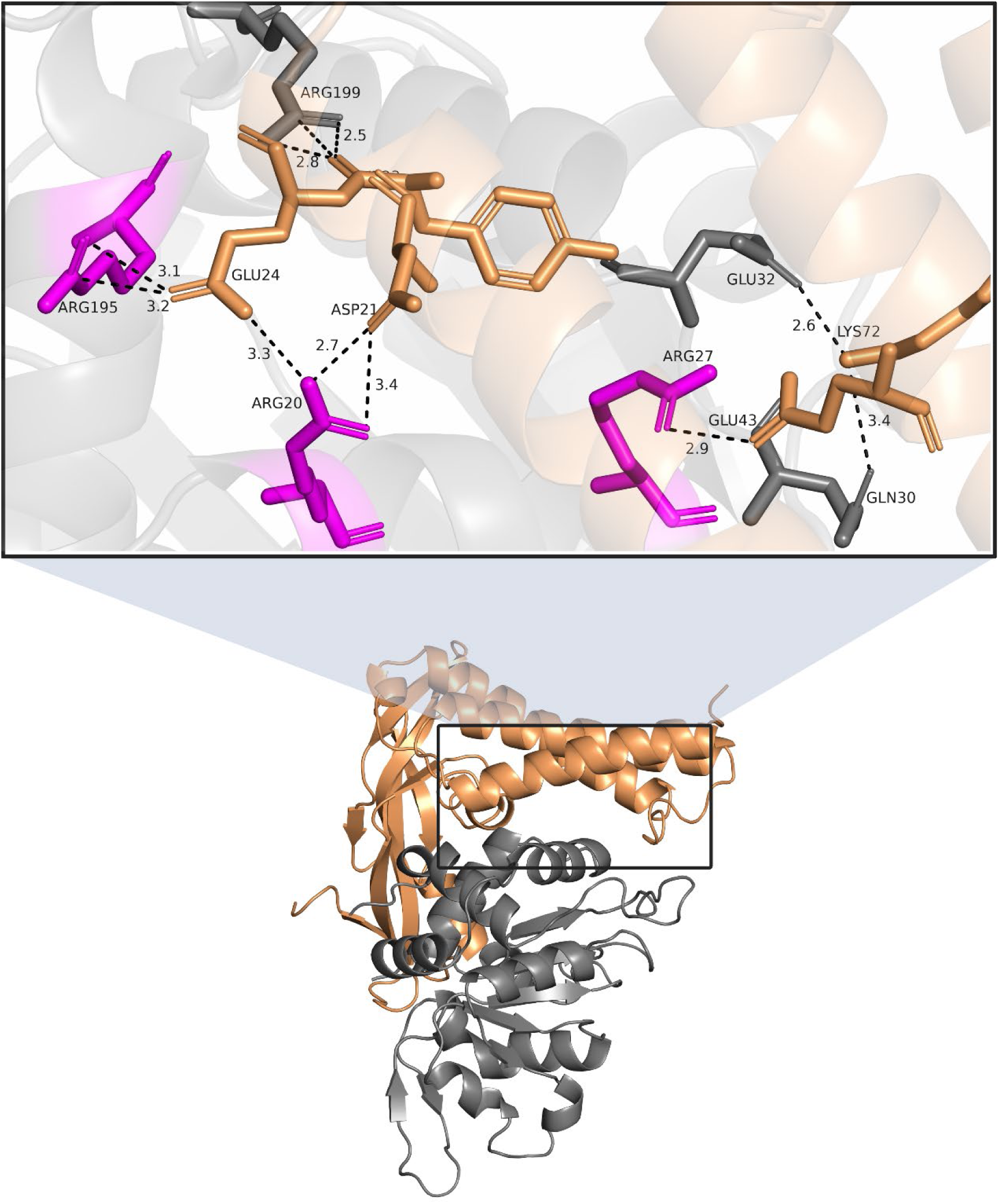
Structural basis of the native HPV-18 E1 and E2 interaction interface. The bottom panel displays the global structure of the E1 helicase domain in grey bound to the E2 transactivation domain in orange. The top panel provides a magnified view of the interaction network. Dashed lines indicate specific interatomic distances in Angstroms between the highly conserved E1 arginine triad residues highlighted in magenta and the complementary acidic residues of the E2 domain.

### 3.2 Toxicity, Allergenicity, and Safety Profiling

All high confidence leads successfully passed the safety assessments by falling well below the strict CSM-Toxin classification threshold of 0.968. **(Fig. 3A)** Scores within the candidate pool ranged from 0.54 for Binder 6 to an optimal score of 0.0 for Binder 8, confirming the general non-toxic nature of the designed sequences. The AlgPred 2.0 systemic allergenicity analysis produced similar findings. While Binders 13 had predictive score of 0.56, which slightly exceeds the standard 0.5 threshold, it was retained for initial baseline characterization with this caveat, while all other binders fell strictly below 0.5. **(Fig. 3B)**

**Fig. 3.**
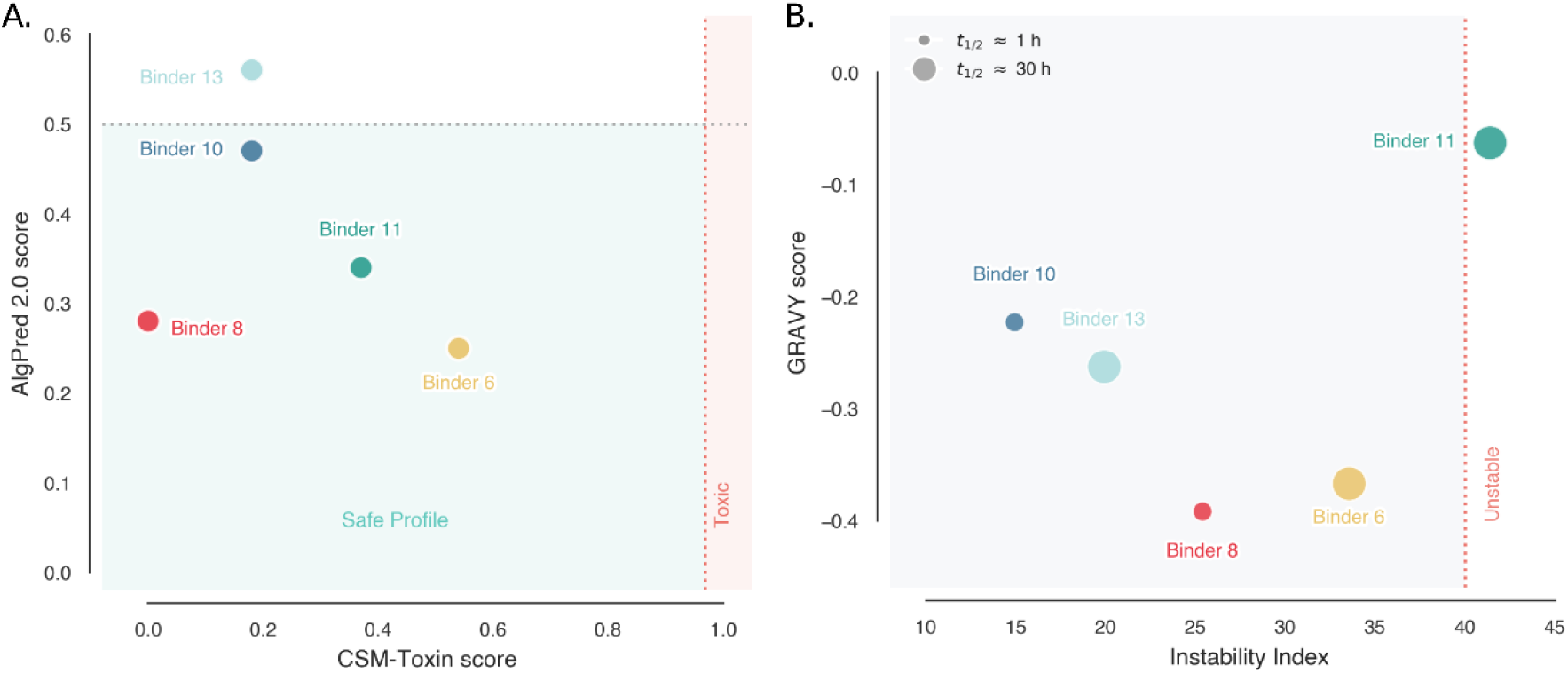
In silico safety and physicochemical profiling of E1-targeted peptide candidates. (A) Cytotoxicity and allergenicity risk assessments evaluated via the CSM-Toxin and AlgPred 2.0 predictive architectures. (B) Physicochemical stability and predicted aqueous solubility derived from ExPASy ProtParam analysis.

### 3.3 Physicochemical Stability and Solubility

ProtParam analysis was used to determine the behavior of the designed peptides in a physiological environment. Binders 6, 8, 10, and 13 are classified as stable with instability index values falling below the standard threshold of 40. Binder 11 yielded an instability index of 41.39, placing it marginally above the stability threshold. The peptides yielded negative GRAVY scores, which confirms high hydrophilicity and predicted aqueous solubility. Binder 8 scored -0.392, Binder 6 scored -0.367, Binder 13 scored -0.263, Binder 10 scored -0.223, and Binder 11 scored -0.063. Additional structural profiling revealed aliphatic indices of 117.92 for Binder 10, 123.67 for Binder 11, and 118.13 for Binder 13. Binders 6, 11, and 13 showed predicted half-lives of 30 hours. Binder 10 showed a predicted half-life of 1.9 hours. Binder 8 displayed a shorter half-life of 1.3 hours due to its N-terminal Lysine.

### 3.4 Thermodynamic Profiling and Mechanistic Energy Decomposition

The thermodynamic evaluation of the AlphaProteo-generated sequences via the dual-scale MD and MM/GBSA protocol yielded distinct binding affinities, successfully isolating 3 strong leads. When balancing highly favorable thermodynamic stability with a superior safety profile, Binder 8 emerged as the overall primary lead candidate. Throughout the statistical sampling, the Binder 8-E1 complex maintained a strong binding free energy (ΔG_MM/GBSA_) of -59.1 ± 0.7 kcal/mol (SEM) with a standard deviation (SD) of 4.95 kcal/mol. **(Fig. 4A)** This represents a highly significant thermodynamic improvement over the native 1TUE reference complex (−48.9 ± 0.6 kcal/mol, SD = 4.24 kcal/mol) (Welch’s t-test, p = 8.14e-19, Bonferroni-corrected). The biological impact of this improved affinity is further supported by a large Cohen’s d effect size of 2.21. Furthermore, to determine the specific residues driving this high binding affinity, per-residue energy decomposition was performed. The quantitative energetic data shows that multiple binders successfully coordinate the native basic residues protruding from the E1 surface. This affinity is anchored by the multi-point interaction network targeting the R20, R27, and R195 Arginine Triad.

**Fig. 4.**
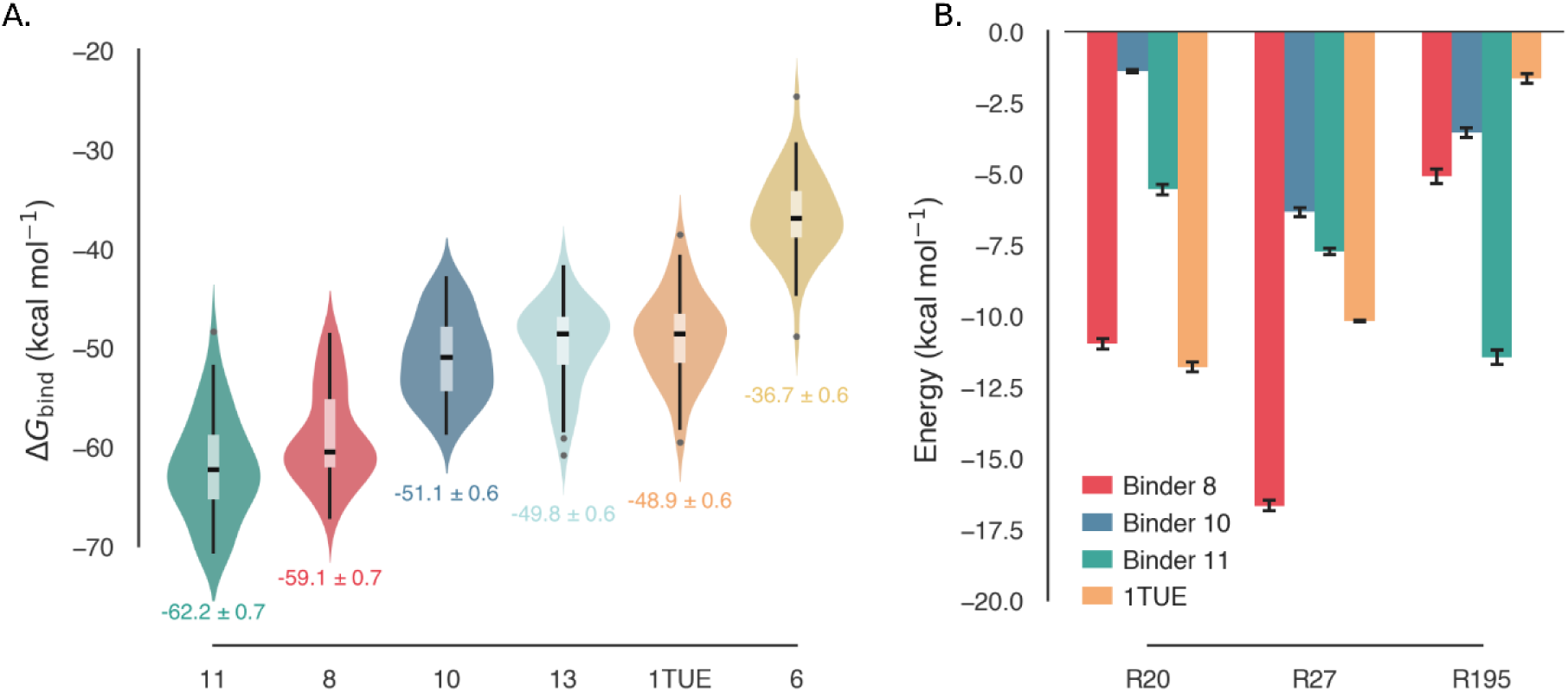
Predicted Binding free energy distributions of the de novo peptide candidates and reference 1TUE. Binding free energies (Molecular Mechanics/Generalized Born Surface Area, ΔG_MM/GBSA_) were calculated across independent replicates. (A) Violin plots detailing the ΔG_MM/GBSA_ distribution profiles for Binders 6, 8, 10, 11, and 13 alongside the native 1TUE reference complex. (B) Targeted per-residue energy decomposition comparing the specific enthalpic contributions of the highly conserved arginine triad residues across Binders 8, 10, and 11 alongside the native 1TUE reference complex.

Residue-specific energetics of Binder 8 reveal that R27 is the primary structural anchor of the complex, contributing -15.23 kcal/mol to the overall binding stability **(Fig. 4B)**. R20 serves as a secondary structural anchor, contributing -8.73 kcal/mol, while R195 supports the periphery of the binding pocket with an affinity of -4.50 kcal/mol. Extended energetic profiling confirms that this triad heavily dominates the interaction profile across all top candidates, with Binder 8 demonstrating the most optimized binding. **(Fig. S2)** Following the thermodynamic evaluation of the initial candidate pool via the MM/GBSA statistical ensemble, Binders 8, 10, and 11 demonstrated the most highly favorable predicted binding free energies. Binder 11 (−62.2 ± 0.7 kcal/mol, SD = 4.95 kcal/mol) and Binder 10 (−51.1 ± 0.6 kcal/mol, SD = 4.24 kcal/mol) both exhibited statistically significant improvements over the native complex (p = 9.52e-26 and p = 1.10e-02, respectively), with Binder 11 demonstrating a similarly large effect size (Cohen’s d = 2.89). **(Table 1)** Consequently, these three sequences were isolated as the lead candidates. To provide structural context for the top performer, Binder 8’s AlphaFold 3 structure is detailed in **Fig. 5**.

**Fig. 5.**
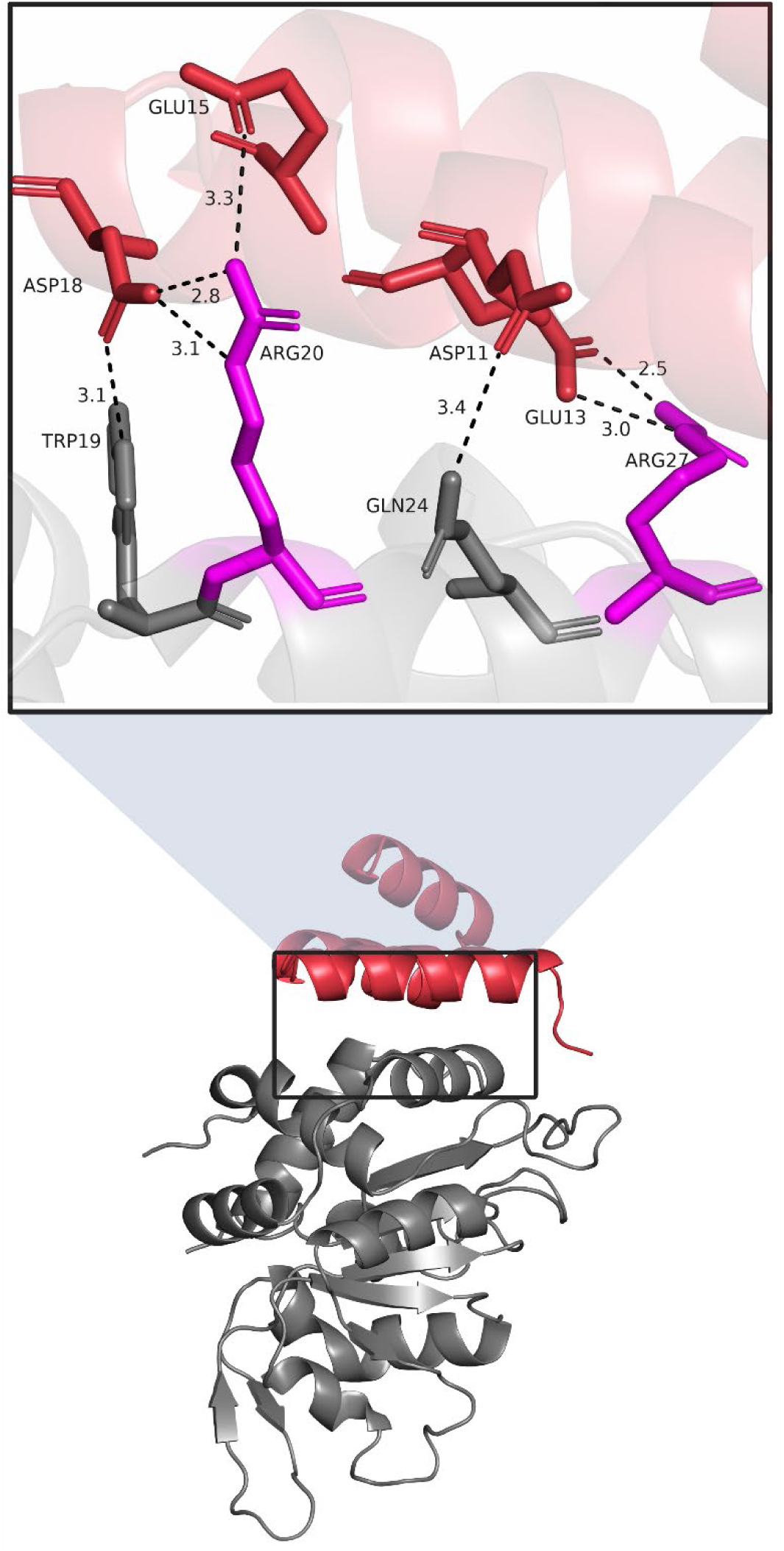
Predicted interaction network of the lead de novo peptide inhibitor Binder 8 complexed with the HPV-18 E1 helicase. The global binding pose of Binder 8 in red to the E1 receptor in grey is shown in the bottom panel. The top panel highlights the atomic interface. The designed peptide is predicted to form a localized electronegative pocket to strongly trap the target E1 arginine residues (R20 and R27). Interatomic distances between key Binder 8 residues and the E1 arginines in magenta are denoted by dashed lines in Angstroms.

**Table 1:** Biophysical and Physicochemical Characterization of Lead Peptide Binders.

| System | $\Delta G_{MM/GBSA}$<br>(kcal/mol) | Toxicity<br>(CSM) | Allergenicity<br>(AlgPred) | Instability<br>Index | GRAVY<br>Score | Predicted<br>Half-life<br>(hrs) |
| --- | --- | --- | --- | --- | --- | --- |
| Binder 8 | $-59.1 \pm 0.7$ | 0.00 | 0.28 | 25.39 | -0.392 | 1.3 |
| Binder 10 | $-51.1 \pm 0.6$ | 0.18 | 0.47 | 14.94 | -0.223 | 1.9 |
| Binder 11 | $-62.2 \pm 0.7$ | 0.37 | 0.34 | 41.39 | -0.063 | 30.0 |
| 1TUE (Native) | $-48.9 \pm 0.6$ | — | — | — | — | — |

Comparative metrics of the top computationally designed peptide inhibitors against the native viral complex (1TUE). The table details
thermodynamic scoring via MM/GBSA, predicted physicochemical properties and deep-learning-based toxicological assessments.
Abbreviations: MM/GBSA, Molecular Mechanics/Generalized Born Surface Area; CSM, CSM-Toxin; AlgPred, AlgPred 2.0; GRAVY, Grand Average of Hydropathy.

### 3.5 Structural Stability

Having established both favorable docking poses and strong predicted binding affinities, these complexes were advanced to the final stability analysis phase. To test the persistence of these interactions under physiological conditions, Triplicate 50 ns long-run trajectory analyses were used to assess the stability of the bound state for the viable candidates (Binders 8, 10, 11), with the RMSD demonstrating a steady, restrained equilibration that remains well below 0.25 nm. These results are significantly more stable than the volatile 1TUE reference, showing no signs of global unfolding or complex dissociation.

Furthermore, independent structural metrics of the isolated peptide binders confirmed their high internal stability **(Fig. S3)**. The Rg of the shared E1 receptor domain remained constant across all evaluated replicates, representing that the structural compactness of the receptor fold is maintained regardless of the bound ligand. Interfacial hydrogen bond analysis, restricted to receptor-ligand contacts only, revealed that the 1TUE complex maintains a mean of 7.7 ± 1.8 interfacial H-bonds, compared to 6.4 ± 1.6 for Binder 11, 4.7 ± 1.5 for Binder 10, and 4.1 ± 1.1 for Binder 8. **(Fig. 6)**

**Fig. 6.**
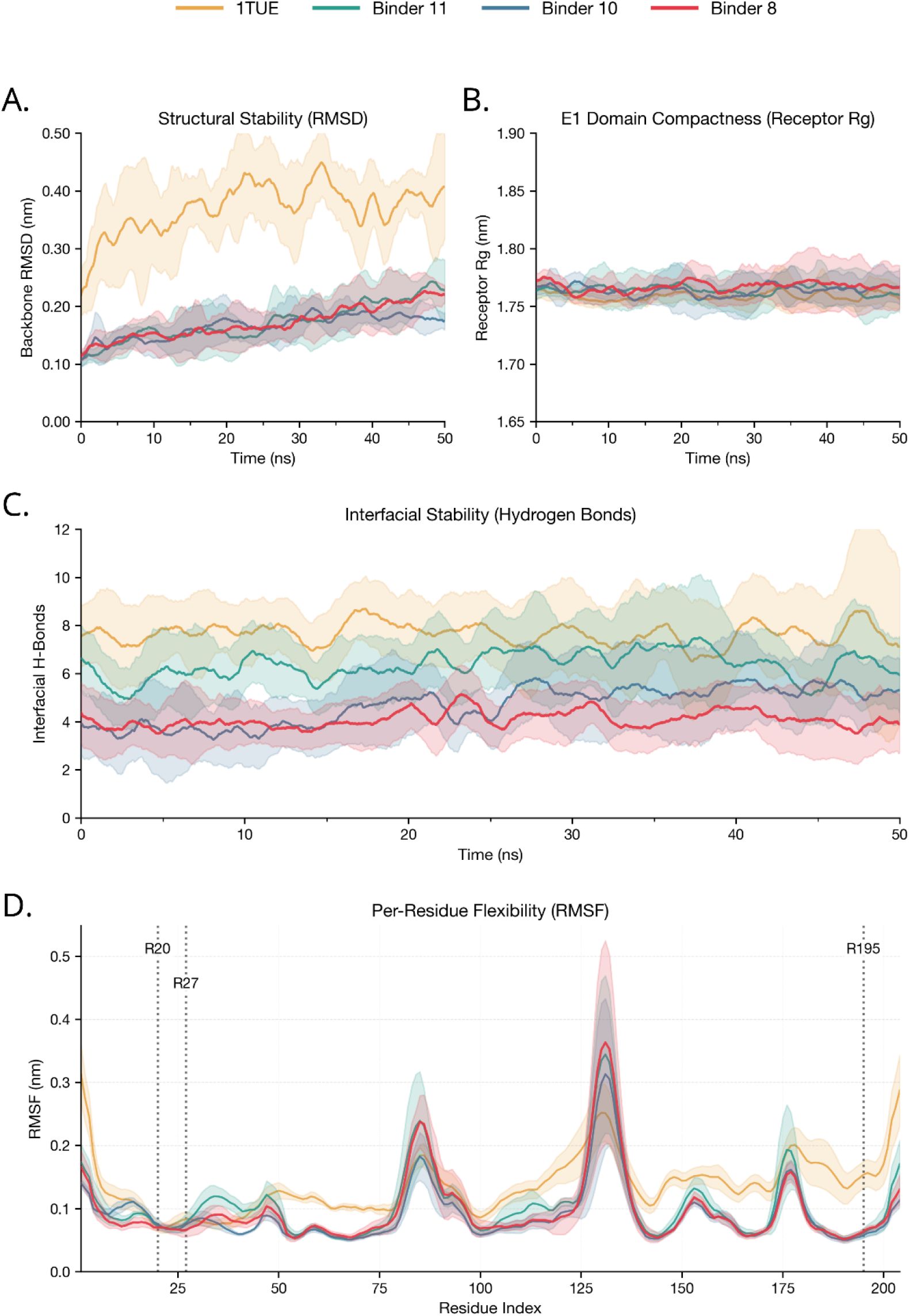
Structural stability of the top 3 Receptor-Peptide complexes and 1TUE during 50 ns molecular dynamics simulations. (A) Root Mean Square Deviation profiles mapping backbone structural variance over time. (B) Radius of Gyration computed over the shared E1 receptor domain (204 residues) to ensure size-independent comparability. (C) Interfacial hydrogen bond counts restricted to receptor-ligand contacts only, excluding intra-chain H-bonds. (D) Per-residue Root Mean Square Fluctuation mapping the local flexibility of the E1 receptor, with the targeted Arginine Triad residues (R20, R27, R195) marked by dotted lines. Note: Faded regions represent standard deviations of triplicate simulations

### 3.6 DCCM analysis

The 1TUE complex (419 C-alpha atoms; 87,571 possible pairs) exhibits a correlated fraction of 21.5% and an anti-correlated fraction of 15.1%. In contrast, Binder 8 (253 C-alpha atoms; 31,878 possible pairs) shows a higher correlated fraction of 25.8% and anti-correlated fraction of 24.9%. Binder 11 yields 20.3% correlated and 13.9% anti-correlated, while Binder 10 shows the most constrained dynamics at 6.6% correlated and 1.2% anti-correlated. Notably, Binder 8 concentrates a greater proportion of its residue pairs into tightly coordinated networks relative to 1TUE. **(Fig. 7).**

**Fig. 7.**
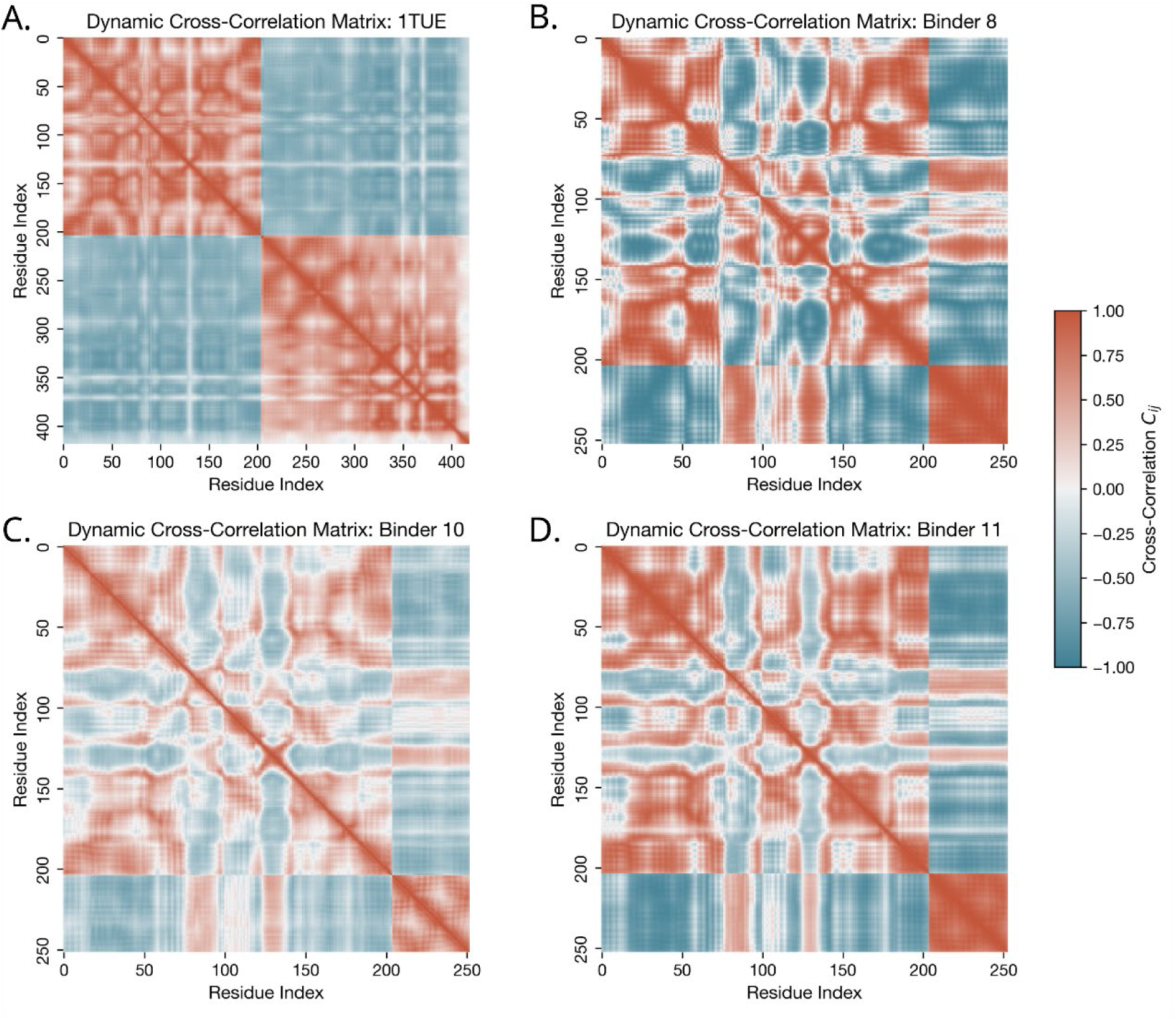
Global conformational dynamics and allosteric network analysis of the E1 protein complexes. Dynamic Cross-Correlation Matrix (DCCM) analyses were executed across the triplicate 50 ns production trajectories. Correlated and anti-correlated pair counts at the |r| > 0.6 threshold are reported as fractions of the total possible unique pairs (N(N-1)/2) to normalize for system size. (A-D) DCCM plots mapping internal residue correlations for the native 1TUE complex (A), Binder 11 (B), Binder 10 (C), and the Binder 8 (D).

### 3.7 DRN and Betweenness Centrality

In the native E1-E2 interaction (1TUE), internal communication is centralized through residues 93 (BC = 0.111), 63 (BC = 0.101), 70 (BC = 0.089), and 100 (BC = 0.072), with average shortest path lengths ranging from 2.6 to 2.9. Binder 8 exhibits a comparable centrality profile anchored by residues 93 (BC = 0.107), 70 (BC = 0.092), 63 (BC = 0.081), and 103 (BC = 0.063), with average shortest path lengths in the range of 2.6 to 3.0. Binder 10 is distinguished by a shift in centrality values, with residue 63 (BC = 0.099) showing a slightly higher score than residue 93 (BC = 0.092). **(Fig. 8)**

**Fig. 8.**
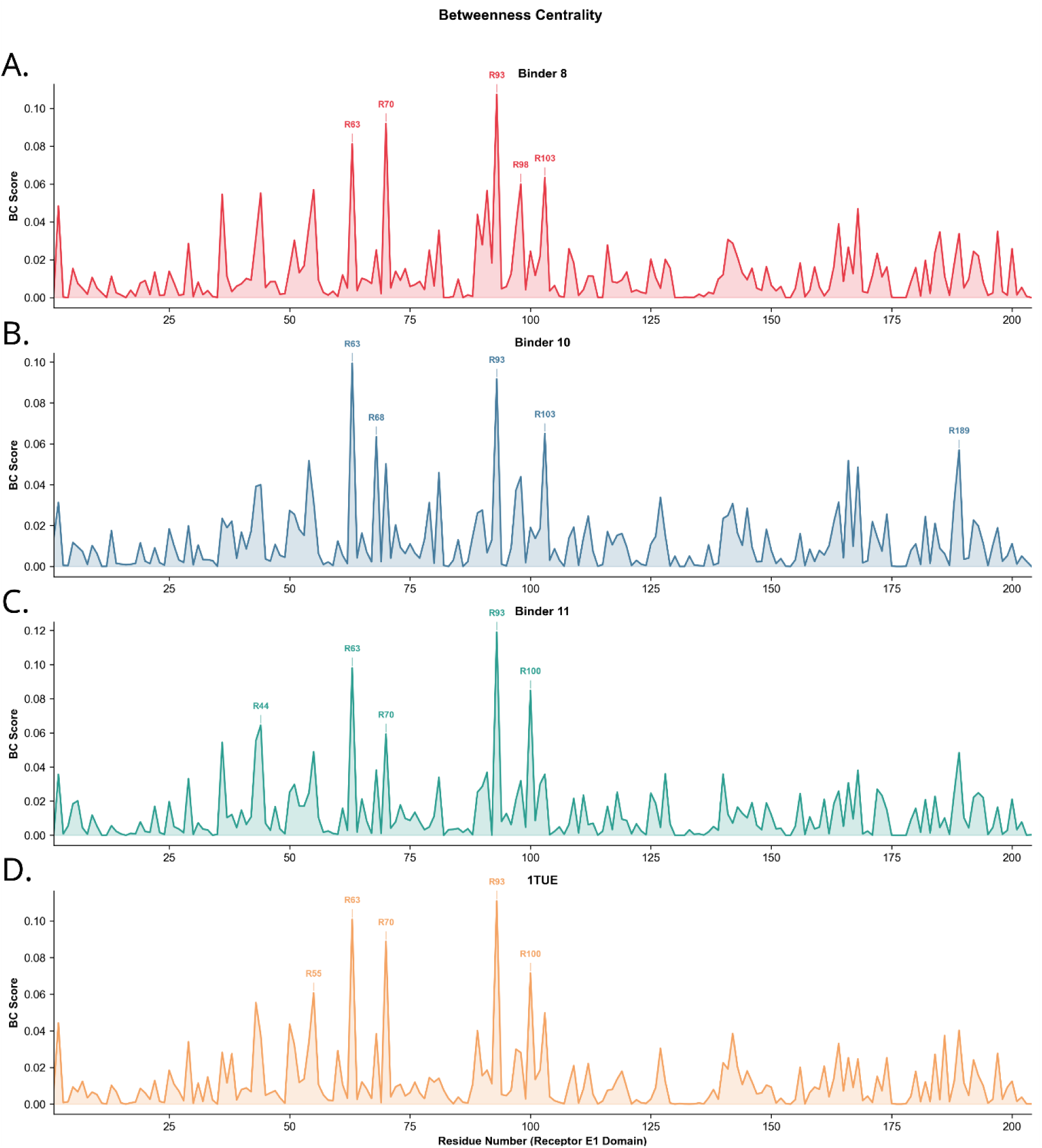
Dynamic residue network analysis of the top 3 binders and reference 1TUE. Betweenness centrality scores were computed exclusively on the shared 204-residue E1 receptor subgraph to ensure identical normalization denominators across systems of different sizes. BC profiles are mapped for: (A) Binder 8, (B) Binder 10, (C) Binder 11, alongside the (D) native 1TUE reference complex.

### 3.8 Evolutionary Conservation

Protein sequence alignment of the oncogenic Alpha-papillomavirus genus (n=183) demonstrated strong basic charge conservation across the target E1 interface. Position R27 exhibited a strict Arginine conservation of 97.81% and a physicochemical (Arginine/Lysine) conservation of 100.00%. Position R20 demonstrated 52.46% strict Arginine conservation and 100.00% physicochemical conservation. Position R195 showed 76.50% strict Arginine conservation and 88.52% physicochemical conservation. Analysis of the extended Alpha, Beta, and Gamma evolutionary baseline (n=194) yielded strict Arginine conservation rates of 15.98%, 67.01%, and 57.73% for positions R20, R27, and R195, respectively. When accounting for physicochemical conservation, the basic motif was maintained at rates of 74.23% for R20, 99.48% for R27, and 97.42% for R195.

## 4. Discussion

The design and validation process of Binder 8 emphasizes the efficiency of integrating generative artificial intelligence with physics-based validation. By synthesizing AlphaProteo sequence generation with dual scale MD, this workflow successfully identified a series of highly promising candidates from a small initial pool of only 14 candidates. The strength of these candidates is structurally rooted in how the inhibitor engages its target. In the native E1-E2 interaction, the E2 transactivation domain utilizes an array of carefully positioned acidic residues to satisfy the geometric constraints of the E1 arginine triad. [7] AlphaProteo’s generative design for Binder 8 successfully replicated this native acidic engagement. The per-residue energy decomposition confirming less than -15 kcal/mol from R27 demonstrates that Binder 8 likely creates a highly localized electronegative binding pocket that captures the positive charges of the triad, stripping away the surrounding hydration shell. This energetic reliance on R27 is highly biologically significant, as it aligns with previous mutational analyses identifying this exact residue as the primary structural feature of the native E1-E2 interaction. [7] In addition to occluding the E2 binding site, our data suggests that Binder 8 alters the allosteric dynamics of the E1 receptor. Size-normalized DCCM results indicate that the designed binders redistribute rather than fully eliminate allosteric motions. Specifically, Binder 8 concentrates a larger fraction of residue-pair correlations (25.8%) into coordinated networks compared to the native 1TUE complex (21.5%). Furthermore, DRN analysis indicates that Binder 8 narrows these internal communication pathways into a more localized topology. This restricted dynamic profile may disrupt the conformational flexibility required for functional hexamerization and support its mechanism as an inhibitor.

Furthermore, the translation of in silico computational discoveries to clinical practice is most frequently halted not by a lack of target efficacy, but by in vivo toxicity. [45,46] In biologic therapeutic development, non-specific off-target binding, membrane perturbation, and immunogenic anaphylaxis are the primary drivers of Phase I clinical trial failures. [47,48] It is within this context that Binder 8’s CSM-Toxin score (0.0) and non-allergenic AlgPred 2.0 classification elevate it from a theoretical academic exercise to a highly promising candidate for subsequent in vitro validation. While the safety profile of Binder 8 positions it as a strong lead, its predicted half-life of 1.3 hours presents a pharmacokinetic challenge. However, this can be readily addressed during future synthesis, standard modifications like N-terminal acetylation are expected to neutralize the N-end rule degradation and extend the serum half-life of Binder 8 for therapeutic use. [49] Moreover, while Binder 11 technically crossed the predictive stability threshold with an index of 41.39, it remains a highly valuable lead. It exhibited the strongest predicted binding affinity of the entire candidate pool at -62.2 kcal/mol. Because its instability is borderline, Binder 11 serves as a promising scaffold for standard chemical optimizations, such as the aforementioned N-terminal acetylation or peptide stapling. These routine modifications could readily resolve its marginal stability deficit while preserving its thermodynamic potency. [49–53]

By successfully designing a protein sequence devoid of known toxicological motifs, Binder 8 minimizes the physiological and financial risks associated with progressing to in vivo murine and non-human primate models. [18] Beyond establishing this safety baseline, a clinical therapeutic must also overcome the genetic variability of the pathogen. A fundamental challenge in the development of broad-spectrum antivirals is the high degree of genotypic diversity among viral strains. [45,46,54] However, the physicochemical conservation of the arginine triad across nearly all HPV types supports Binder 8’s clinical utility. Currently, treatments for CIN or invasive oropharyngeal cancers are largely surgical or involve highly destructive, broad-spectrum chemoradiation, lacking any true viral specificity. [55–59] A therapeutic agent capable of arresting episomal viral replication across multiple high-risk genotypes could offer a valuable non-surgical intervention for early-stage neoplastic lesions, significantly advancing the clinical management of HPV. [60,61]

Even with the advanced integration of generative AI and classical molecular mechanics, the predictive nature of this computational research requires the honest acknowledgment of fundamental methodological limitations. A primary limitation of this in silico framework is the reliance on the proprietary AlphaProteo architecture for the initial generative design phase. Because the model weights are not currently accessible to the public, direct replication of the specific de novo sequence generation step is restricted. However, the biophysical evaluation pipeline established in this study is inherently model agnostic. The overarching workflow can be coupled with readily available open-source generative models. Platforms such as RFdiffusion [62], its subsequent iteration RFdiffusion3 [63], and LigandMPNN [64] offer highly capable open-access alternatives for targeting specific motifs like the E1 Arginine Triad. Furthermore, emerging low-barrier generative web interfaces continue to democratize access to these structural algorithms. [65–68]

Beyond the generative phase, the thermodynamic validation carries inherent limitations. The MM/GBSA methodology utilized for thermodynamic scoring is an implicit solvent model. While it utilizes the Generalized Born approximation to mathematically represent the electrostatic screening of water, it inherently strips away discrete, explicit water molecules during the energy calculation. In complex PPIs, structural bridging water molecules frequently play a role in stabilizing highly specific interfacial geometries. The absence of explicit solvent networks in the final ΔG_MM/GBSA_ calculation can lead to the systemic overestimation of absolute binding affinities, though the relative thermodynamic ranking of binders (e.g., the superiority of Binder 8 over Binder 6) remains highly reliable and accurate. [36,69,70] Additionally, standard MM/GBSA calculations frequently omit the calculation of configurational entropy (the thermodynamic loss of structural flexibility upon binding) due to the prohibitive computational cost and the known inaccuracies of the rigid-rotor harmonic oscillator approximation for large, flexible peptides. Consequently, the reported -59.1 kcal/mol represents an effective binding enthalpy heavily favored by optimized electrostatic interactions, rather than a true absolute free energy incorporating all entropic penalties.

Furthermore, while advanced deep-learning tools like CSM-Toxin provide high-confidence safety predictions, they are strictly bounded by the inherent biases and limitations of their training datasets. [18] Similarly, Physical pharmacokinetic properties, such as precise in vivo proteolytic degradation rates in serum, physical clearance by the reticuloendothelial system, and deep tissue biodistribution, cannot be fully captured by the ProtParam instability index or primary sequence-based half-life estimations. Finally, A critical translational hurdle for Binder 8 is its requisite intracellular delivery. Because the HPV E1 helicase functions within the nucleus, a biologic peptide must successfully traverse both the cellular and nuclear membranes. Traditional unprotected peptides frequently suffer from poor cellular uptake and endosomal entrapment. [71–73] However, established delivery strategies have successfully propelled similar biologic inhibitors targeting intracellular and nuclear protein-protein interactions into advanced clinical trials. These successful candidates overcome membrane barriers by utilizing advanced delivery modalities, such as macrocyclization or direct conjugation to Cell-Penetrating Peptides. [48,73–77] Future formulations of Binder 8 will need to explore these validated modalities to ensure the protein achieves functionally relevant concentrations at the intracellular target site. Finally, translating these in silico findings into clinical therapeutics will involve transitioning this framework into in vitro models. Future efforts will logically progress toward the empirical validation of Binder 8 by utilizing Bio-layer interferometry (BLI) to confirm the true equilibrium dissociation constant of the engineered binders.

## 5. Conclusion

The integration of generative deep learning models with MD has successfully yielded a series of highly promising, de novo peptide inhibitors targeting the fundamental replication machinery of oncogenic Human Papillomavirus. By explicitly aiming for the conserved triad located on the solvent-exposed surface of the E1 protein, Binder 8 achieves an highly favorable predicted binding free energy (−59.1 kcal/mol) while retaining a safe physicochemical profile, as validated by deep-learning toxicity and allergenicity screens.

Because this interface is near-universally conserved across all HPV strains, Binder 8 represents a highly viable, broad-spectrum antiviral biologic candidate capable of disrupting the E1-E2 pre-initiation complex. This research provides a computationally validated blueprint that directly sets the stage for in vitro BLI kinetics testing, offering a new trajectory in the pharmacological eradication of HPV-driven malignancies.

## Supporting information

Graphica, abstract

## Data Availability Statement

To ensure computational reproducibility and scientific transparency, the complete computational pipeline and structural models used in this study have been deposited in a persistent Zenodo repository (https://doi.org/10.5281/zenodo.20077263). This archive includes the AlphaProteo generation parameters, AlphaFold 3 structural models (.cif format), and the full suite of GROMACS simulation and post-processing scripts required to reproduce the methodology reported in the manuscript.

## CRediT Author Statement

**Sean Fletcher:** Conceptualization, Formal analysis, Methodology, Visualization, Writing - original draft, Writing - review and editing.

**Esther E. Biswas-Fiss:** Conceptualization, Supervision, Funding acquisition.

**Subhasis B. Biswas:** Conceptualization, Methodology, Supervision, Resources, Writing - review and editing, Funding acquisition.

## Ethics Statement

This research is entirely computational and conducted strictly in silico. Therefore, this study did not involve human subjects or animal experiments and is exempt from ethics approval.

## Funding

This work was supported by the National Institute of General Medical Sciences of the National Institutes of Health [grant number P20 GM103446], the State of Delaware, the Delaware INBRE Summer Research Program, and the Angela Santoro Summer Research Award.

## Acknowledgments

The authors thank the members of the Biswas Laboratory for their help and support. Furthermore, we acknowledge and extend our deepest thanks to the Google DeepMind AlphaProteo trusted tester program for officially granting us computational access to their generative AI tool, without which the sophisticated de novo sequence design utilized in this study would not have been possible.

## Declaration of Generative AI and AI-Assisted Technologies in the Manuscript Preparation Process

During the conduct of this research, the authors used AlphaProteo (Google DeepMind) for the de novo generation of candidate peptide binder sequences targeting the HPV E1 helicase arginine triad, as described in Section 2.1. AlphaFold 3 (Google DeepMind) was used for three-dimensional structural prediction of the resulting receptor-peptide complexes, also as described in Section 2.1. These tools were used exclusively as research instruments within the described methodology. All AI-generated outputs, including candidate sequences and structural models, were reviewed, validated, and interpreted by the authors using independent computational methods (Molecular Dynamics simulation, MM/GBSA thermodynamic analysis, and PyMOL structural visualization). The authors reviewed and edited all content and take full responsibility for the scientific conclusions of this article.

## Supplementary Material

**Fig. S1.** Structural superimposition of the AlphaFold 3 predicted HPV-18 E1-E2 complex and the native crystal structure. The AlphaFold 3 predicted model (orange) is aligned to the native E1-E2 crystal structure (grey, PDB: 1TUE). The structural alignment yields an all-atom Root Mean Square Deviation (RMSD) of 0.674 Å, validating the accuracy of the computational prediction baseline.

**Fig. S2.** Extended per-residue energy decomposition. An expanded thermodynamic breakdown detailing the top 15 enthalpic contributions to the binding free energy from the E1 receptor protein

**Fig. S3.** Structural stability metrics of the isolated peptide binders. A. Root Mean Square Deviation and B. Root Mean Square Fluctuation profiles detailing the internal stability and local flexibility of Binders 8, 10, and 11 during the triplicate 50 ns molecular dynamics simulations.

**Table S1.** De novo peptide inhibitor sequences. A list detailing the primary amino acid sequences for all 14 computationally generated candidate binders evaluated in this study. **Table S2. Analyzed HPV genotypes.** A complete list of the high-risk and low-risk HPV strains included in the multiple sequence alignments to determine the evolutionary conservation of the E1 arginine triad.

## Notes

### Competing Interest Statement

The authors have declared no competing interest.

