## Supplementary files for "De Novo Design and Computational Validation of a High-Affinity Peptide Inhibitor Targeting the HPV E1-E2 Interface"

| **Binder ID** | **Amino Acid Sequence** |
|---|---|
| Binder 1 | MTREEIIERLQALGYSRETAERIYRTAVELTGSEEGAIELLEEYIESLT |
| Binder 2 | SLRDLLDAIGDRFLENPDALDPELLRALMDAIEAGDYERATALAEKLLSQ |
| Binder 3 | ETAEETAELLLFAHDGDKEAALRAIEVGVELGVFTEEEAQQVREAIDKLA |
| Binder 4 | MSERERDAEEALRLILEACGYITEDEILAAAKEYELDPETVARIAERLMA |
| Binder 5 | SLEELKKELKEAIKANDLDKVIDISFKILDETGSFEKAGKLIEEAIKEVK |
| Binder 6 | MTLDERIEELAKKYNVSVDTVFKIVTDAEIDKGLSDEEAIEYAEKILSK |
| Binder 7 | MDPERLARALDAARILLEAGVPREEIVASLCRHFDLSEETAQAIVDEAEK |
| Binder 8 | KPDPKVEELADLECELLDRGLWEELERVNRLVEAGKVDEAIAYARAVLE |
| Binder 9 | RPSLEELRELARDLMECGCSDEEIIAFLQSEGLSEEEARRIVAEVRAELA |
| Binder 10 | SLEERIKETAEKYGISVEELTRAVNLARELGFEVDDDLIDLIAEALNS |
| Binder 11 | GLSVEQARELALELHREKLATLEEIRRGLIAAGLSRETAEAIVEEVEAA |
| Binder 12 | SVREDAAKINELILANKIEEATELAKETMAKYGIDAVELMELIGEVAAE |
| Binder 13 | MDLEEAVELAKLLLEAGYSDEEIIKELVKAGLDRETAEKALAEAKKEL |
| Binder 14 | LSEEEVLDLAILLLEAGASEEEVIASLEAAGVSRETAERAVREARAILAR |

| | | | | | | | |
|---|---|---|---|---|---|---|---|
| HPV2 | HPV27 | HPV54 | HPV83 | HPV108 | HPV133 | HPV160 | HPV185 |
| HPV3 | HPV28 | HPV56 | HPV84 | HPV109 | HPV134 | HPV161 | HPV186 |
| HPV4 | HPV29 | HPV57 | HPV85 | HPV110 | HPV135 | HPV162 | HPV187 |
| HPV5 | HPV30 | HPV58 | HPV86 | HPV111 | HPV136 | HPV163 | HPV188 |
| HPV6 | HPV31 | HPV59 | HPV87 | HPV112 | HPV137 | HPV164 | HPV189 |
| HPV7 | HPV32 | HPV60 | HPV88 | HPV113 | HPV138 | HPV165 | HPV190 |
| HPV8 | HPV33 | HPV61 | HPV89 | HPV114 | HPV139 | HPV166 | HPV191 |
| HPV9 | HPV34 | HPV62 | HPV90 | HPV115 | HPV140 | HPV167 | HPV192 |
| HPV10 | HPV35 | HPV65 | HPV91 | HPV116 | HPV141 | HPV168 | HPV193 |
| HPV11 | HPV36 | HPV66 | HPV92 | HPV117 | HPV142 | HPV169 | HPV194 |
| HPV12 | HPV37 | HPV67 | HPV93 | HPV118 | HPV143 | HPV170 | HPV195 |
| HPV13 | HPV38 | HPV68 | HPV94 | HPV119 | HPV144 | HPV171 | HPV196 |
| HPV14 | HPV39 | HPV69 | HPV95 | HPV120 | HPV145 | HPV172 | HPV197 |
| HPV15 | HPV40 | HPV70 | HPV96 | HPV121 | HPV146 | HPV173 | HPV199 |
| HPV16 | HPV42 | HPV71 | HPV97 | HPV122 | HPV147 | HPV174 | HPV200 |
| HPV17 | HPV43 | HPV72 | HPV98 | HPV123 | HPV148 | HPV175 | HPV201 |
| HPV18 | HPV44 | HPV73 | HPV99 | HPV124 | HPV149 | HPV176 | HPV202 |
| HPV19 | HPV45 | HPV74 | HPV100 | HPV125 | HPV150 | HPV177 | HPV228 |
| HPV20 | HPV47 | HPV75 | HPV101 | HPV126 | HPV151 | HPV178 | HPV229 |
| HPV21 | HPV48 | HPV76 | HPV102 | HPV127 | HPV152 | HPV179 | |
| HPV22 | HPV49 | HPV77 | HPV103 | HPV128 | HPV153 | HPV180 | |
| HPV23 | HPV50 | HPV78 | HPV104 | HPV129 | HPV154 | HPV181 | |
| HPV24 | HPV51 | HPV80 | HPV105 | HPV130 | HPV155 | HPV182 | |
| HPV25 | HPV52 | HPV81 | HPV106 | HPV131 | HPV156 | HPV183 | |
| HPV26 | HPV53 | HPV82 | HPV107 | HPV132 | HPV159 | HPV184 | |

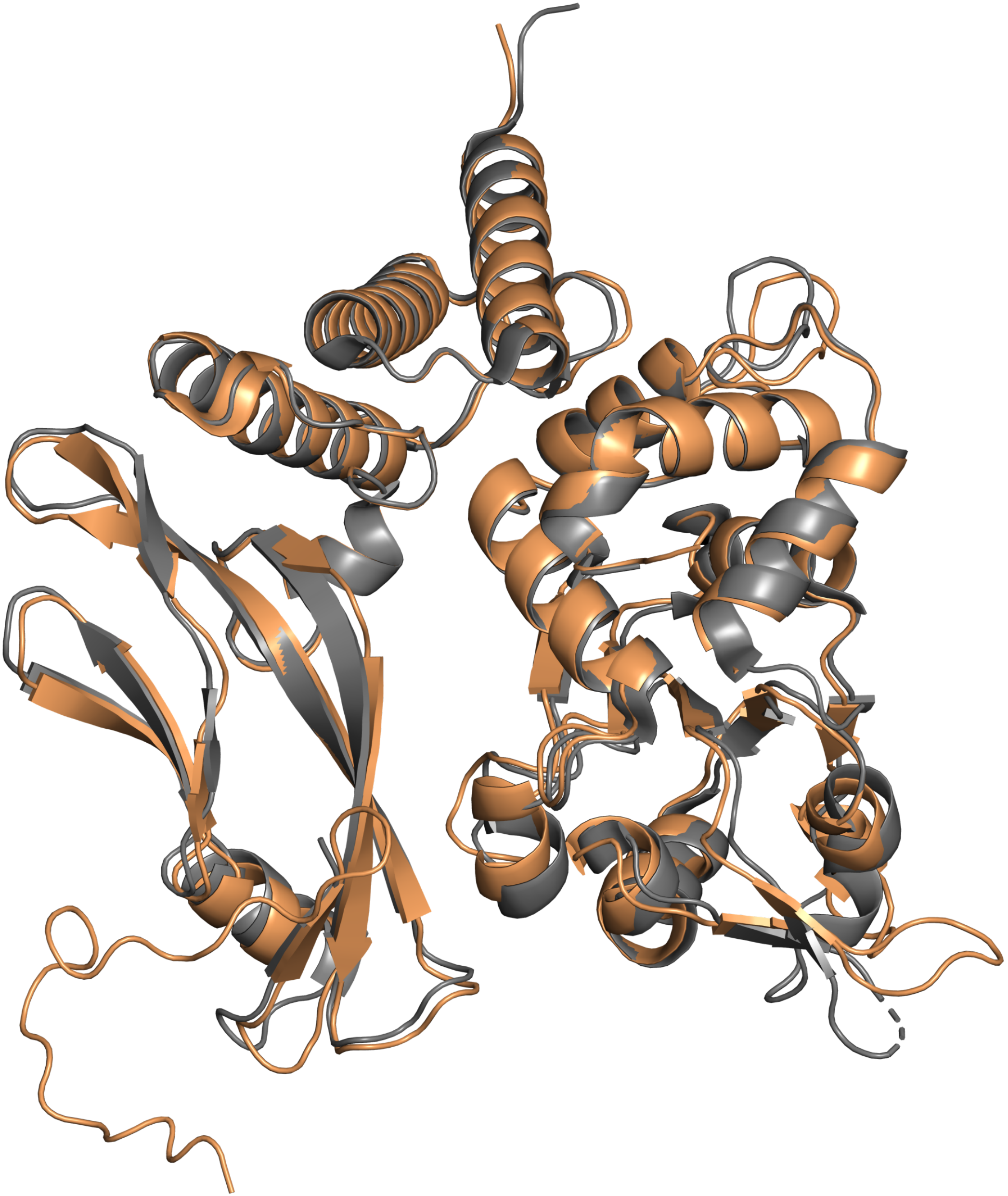

RMSD: 0.674 Å

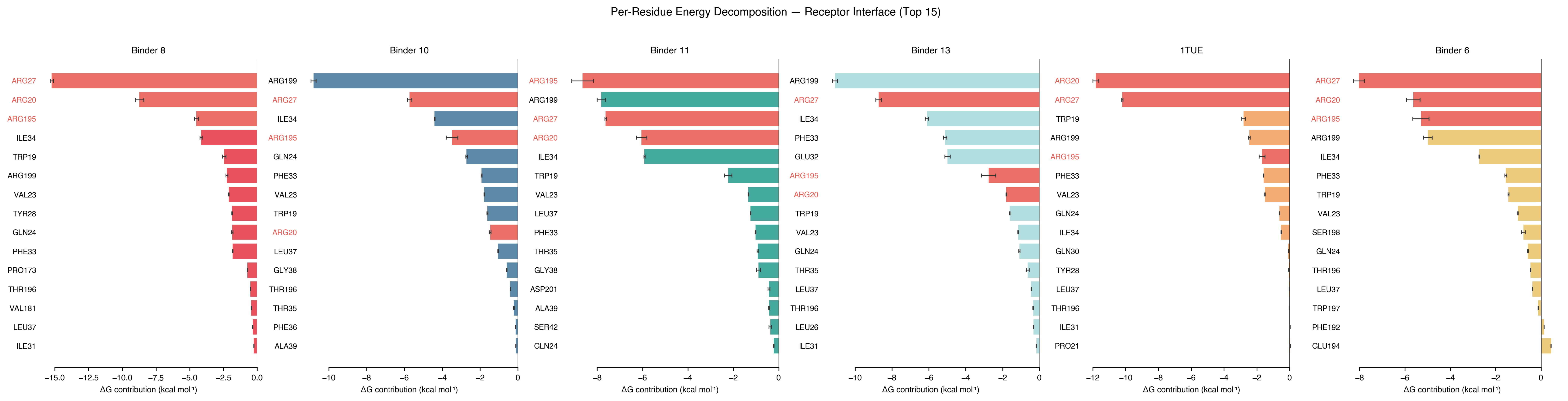

Per-Residue Energy Decomposition — Receptor Interface (Top 15)
Binder 8
ARG27
ARG20
ARG195
ILE34
TRP19
ARG199
VAL23
TYR28
GLN24
PHE33
PRO173
THR196
VAL181
LEU37
ILE31
−15.0
−12.5
−10.0
−7.5
−5.0
−2.5
0.0
ΔG contribution (kcal mol⁻¹)
Binder 10
ARG199
ARG27
ILE34
ARG195
GLN24
PHE33
VAL23
TRP19
ARG20
LEU37
GLY38
THR196
THR35
PHE36
ALA39
−10
−8
−6
−4
−2
0
ΔG contribution (kcal mol⁻¹)
Binder 11
ARG195
ARG199
ARG27
ARG20
ILE34
TRP19
VAL23
LEU37
PHE33
THR35
GLY38
ASP201
ALA39
SER42
GLN24
−8
−6
−4
−2
0
ΔG contribution (kcal mol⁻¹)
Binder 13
ARG199
ARG27
ILE34
PHE33
GLU32
ARG195
ARG20
TRP19
VAL23
GLN24
THR35
LEU37
THR196
LEU26
ILE31
−10
−8
−6
−4
−2
0
ΔG contribution (kcal mol⁻¹)
1TUE
ARG20
ARG27
TRP19
ARG199
ARG195
PHE33
VAL23
GLN24
ILE34
GLN30
TYR28
LEU37
THR196
ILE31
PRO21
−12
−10
−8
−6
−4
−2
0
ΔG contribution (kcal mol⁻¹)
Binder 6
ARG27
ARG20
ARG195
ARG199
ILE34
PHE33
TRP19
VAL23
SER198
GLN24
THR196
LEU37
TRP197
PHE192
GLU194
−8
−6
−4
−2
0
ΔG contribution (kcal mol⁻¹)

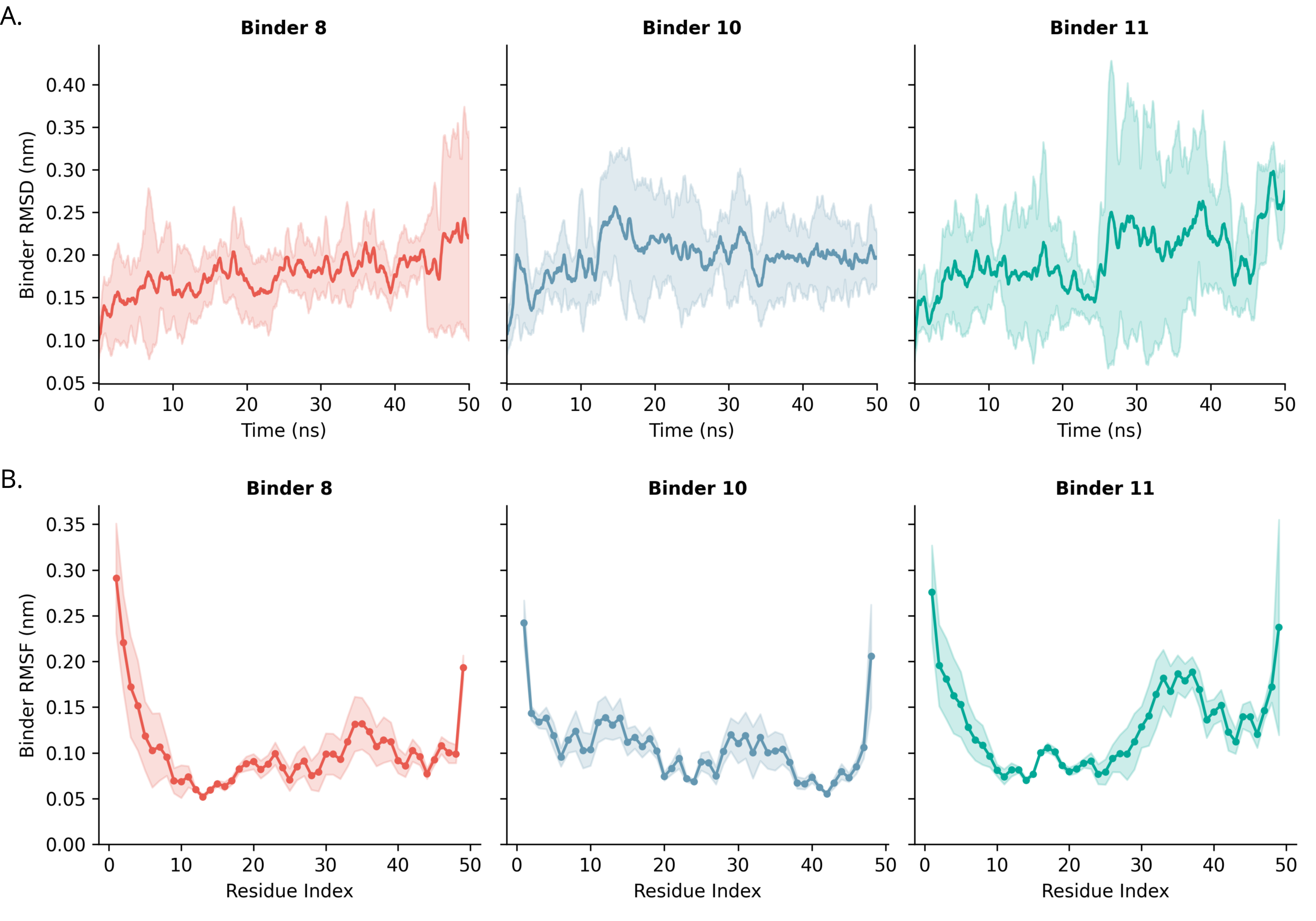

A.
Binder 8
Binder 10
Binder 11
Binder RMSD (nm)
0.05
0.10
0.15
0.20
0.25
0.30
0.35
0.40
0
10
20
30
40
50
Time (ns)
B.
Binder 8
Binder 10
Binder 11
Binder RMSF (nm)
0.00
0.05
0.10
0.15
0.20
0.25
0.30
0.35
Residue Index
