## Supplementary figures and images for "De Novo Design and Computational Validation of a High-Affinity Peptide Inhibitor Targeting the HPV E1-E2 Interface"

### Graphica, abstract

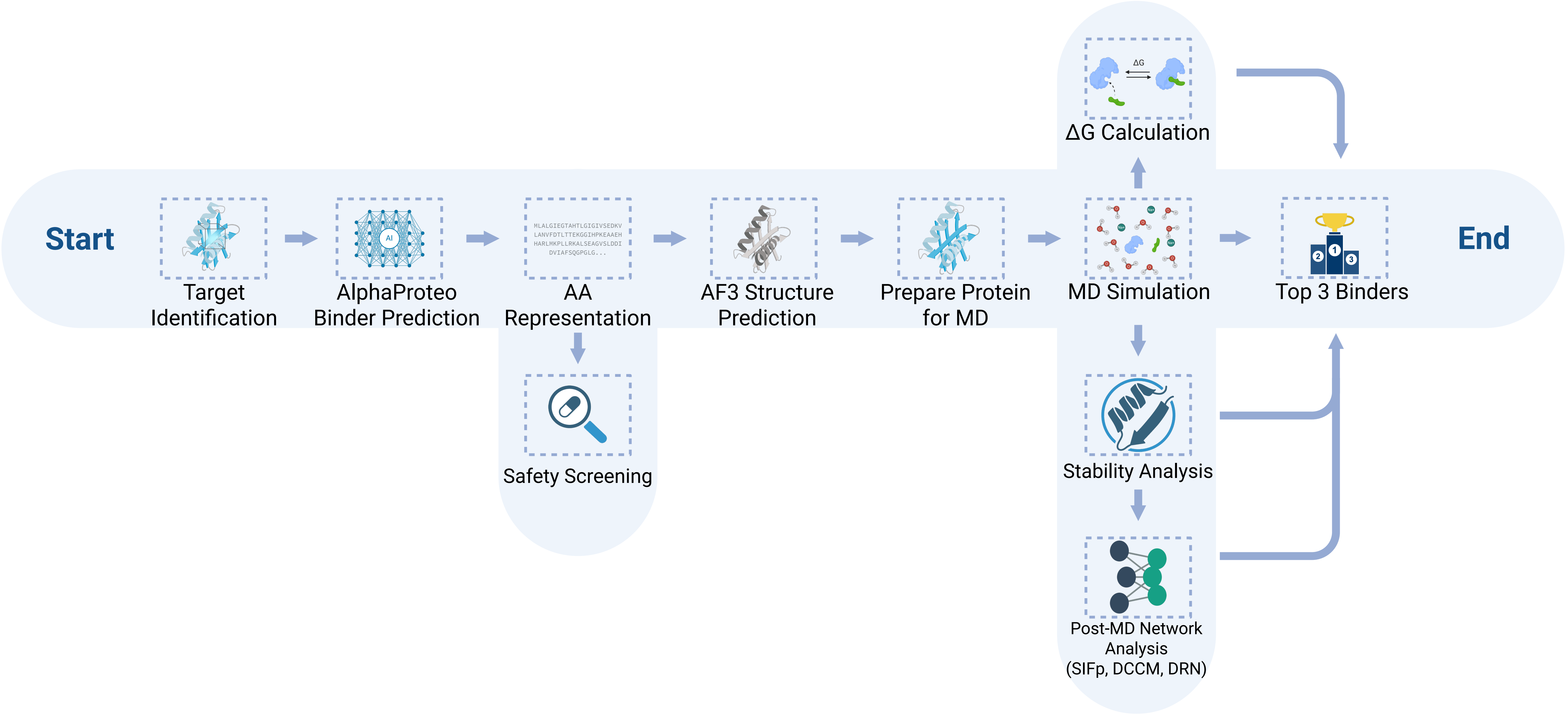
